# A New Generalized Gamma-Weibull Distribution with Applications to Time-to-event Data

**DOI:** 10.1101/2023.11.18.567670

**Authors:** Kazeem Adesina Dauda, Rasheed Kehinde Lamidi, Adeshola Adediran Dauda, Waheed Babatunde Yahya

**Affiliations:** Department of Mathematics, University of Bergen, 5007 Bergen, Norway; Department of Mathematics and Statistics, Kwara State University, Malete, Nigeria; Department of Statistics, University of Ilorin, Nigeria; alt-Address: Department of Mathematics and Statistics, Kwara State University, Malete, Nigeria

**Author notes:** Email addresses:* (Kazeem Adesina Dauda), (Rasheed Kehinde Lamidi), (Adeshola Adediran Dauda), (Waheed Babatunde Yahya).

**Keywords:** generalized Weibull, Gamma-generated distributions, moments, order statistic

## Abstract

In this research, a new class of probability distributions referred to as Generalized Gamma Weibull (GGW) distributions was introduced within the context of parametric survival analysis. This distribution represents a modification of the gamma Weibull distribution and offers valuable insights, particularly when dealing with highly skewed lifetime data. The study extensively examined the mathematical characteristics of these distributions, encompassing hazard functions, moments, quantile functions, and order statistics. Furthermore, the research delved into parameter estimation methods for these newly proposed distributions, employing the maximum likelihood technique, Fisher Information (FI), and deriving asymptotic confidence intervals for both censored and uncensored scenarios. To illustrate the practical utility of these proposed distributions, the study applied them to analyze two sets of real-life survival data and two sets of real-life data, resulting in a total of four distinct datasets. To gauge the effectiveness of the GGW distributions in comparison to existing methods such as Generalized Weibull and Generalized gamma (G-Weibull and G-Gamma) distributions, the research employed statistical indices including the Akaike Information Criterion (AIC), Corrected Akaike Information Criterion (CAIC), and Bayesian Information Criterion (BIC). The outcomes of this comparative analysis demonstrated the superior performance of the newly introduced GGW distributions (AIC=338.6313, BIC=346.2794, and CAIC=339.5202) when contrasted with the existing methods (G-Weibull: AIC=376.1946, BIC=381.9307, and CAIC=376.5424) across all three criteria, thereby highlighting the enhanced suitability of GGW distributions for modeling and analyzing skewed lifetime data.

## 1. Introduction

Zografos and Balakrishnan [1], as well as Ristić and Balakrishnan [2], have introduced two distinct distribution categories, each generated using gamma random variables, which incorporate an additional positive shape parameter. The mathematical characteristics of this distribution were examined by [3], with a specific focus on its applicability to highly skewed time-to-event data, while disregarding the censoring information in the dataset. In separate investigations, [4] and [5] have devised a novel distribution by combining elements of the Zografos-Balakrishnan log-logistic distribution and the gamma exponentiated-Weibull distribution. When compared to the original distribution proposed by ([1]), this new distribution exhibited remarkable performance improvements. Furthermore, [6] has put forth an innovative variation within the Zografos-Balakrishnan family, introducing the Zografos-Balakrishnan-Gompertz distribution and the Zografos-Balakrishnan Lindley distribution. In their work, they meticulously established the mathematical properties, including moments, hazard rate functions, and reliability functions, for these newly introduced distributions. These distributions prove to be highly effective for modeling datasets that exhibit a pronounced skewness. Taking a broader perspective, [6] delved into a generalized form of the log-transformed inverse Weibull distribution, conducting an extensive investigation of its theoretical properties. In summary, several studies in the literature, such as those cited in references ([7, 8]), have explored and developed flexible distribution models capable of handling highly skewed time-to-event data. However, there still remains a need for the development of a distribution that can adeptly accommodate highly skewed time-to-event data while simultaneously considering censoring information.

### 1.1 Existing and Parent Models

The probability density function (pdf) *f* (*y*) and cumulative distribution function (cdf) *F* (*y*) for the distribution family introduced by [1] is expressed as follows:

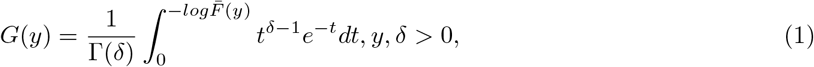

where *F* (*y*)(i.e the cdf) is the parent or based distribution. However, the pdf of the generator is defined as;

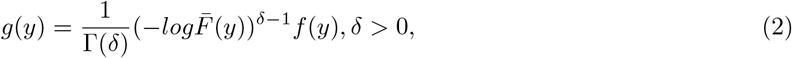

where Γ(.) is the gamma function. The derivation of the new distributions relies solely on these equations. The corresponding Hazard rate function (hrf) is;

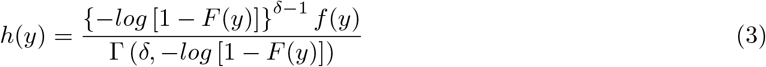

where 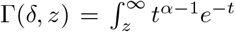 denotes the complementary incomplete gamma function. The generator of the hazard rate function has an additional shape parameter *δ >* 0.

This study considers introducing a new generalized gamma-weibull distribution, inspired by the findings of Zagrafos and Balakrishnan’s distribution family described in equation 1. A new modified generalized gamma-Weibull distribution has been developed as a member of the Zagrofos and Balakrishnan distribution family [1]. A similar version of the generalized gamma-Weibull distribution has been introduced in the master thesis of one of the authors of this study in 2022[9]. In this new version, we derive the statistical properties of this distribution, estimate the maximum likelihood, and provide an application example. Moreover, it is a prevalent practice in the literature to focus on uncensored observations while disregarding censoring scenarios. However, our study addresses these limitations by incorporating more adaptable parameters that accommodate both censored and uncensored observations. Further elaboration on the proposed distributions can be found in the subsequent section.

The structure of the paper unfolds as follows: In Section 2, we express *g*(*y*) in equation 5 as an infinite linear combination of exponentiated gamma-Weibull density functions. Section 2.3 is dedicated to providing the Quantile function for the Generalized Gamma Weibull distribution. Section 2.4 focuses on obtaining moments, while Section 2.5 delves into the distribution of order statistics. Moving forward to Section 2.6, we employ maximum likelihood estimation to determine the model parameters. The Asymptotic confidence interval for the GGW distribution is derived in Section 3. In Section 4, we use censored samples to estimate the model parameters using maximum likelihood in the Generalized Gamma Weibull distribution. The methods of model evaluation is outlined in Section 5. Finally, Section 6 illustrates the practical utility of the approach through its application to a real-life dataset. Concluding remarks are presented in Section 7.

## 2. Methodology

The generation of new distributions relies on the use of the Generalized Weibull (GW) distribution, which was originally proposed by Mudholkar and Kumar [10]. The Generalized Weibull distribution is characterized by its probability density function (PDF) denoted as *f* (*y*) and cumulative distribution function (CDF) represented as *F* (*y*). The PDF and CDF expressions can be found in equations 4 and 5.

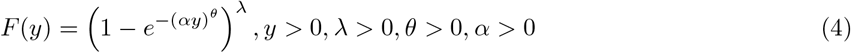

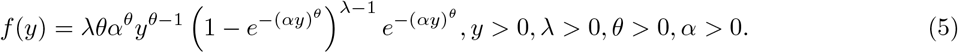

We initiate the creation of this novel distribution by substituting equation 4 into equation 1 and equation 5 into equation 2, respectively. This process results in the cumulative distribution function (CDF) of the newly modified distribution, referred to as the Generalized Gamma Weibull (GGW), which is derived and presented in equation 6 and equation 7.

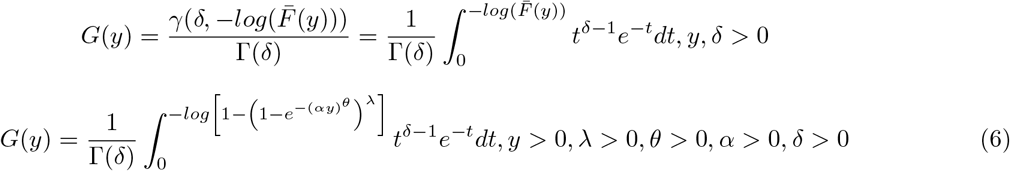

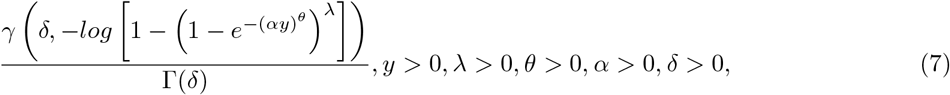

where *γ*(.) is called power series see [11] and define as

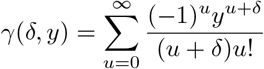

In a similar manner, the probability density function (PDF) of the Generalized Gamma Weibull (GGW) is acquired by substituting the quantity from equation 5 into equation 2. The resulting PDF is then expressed as follows:

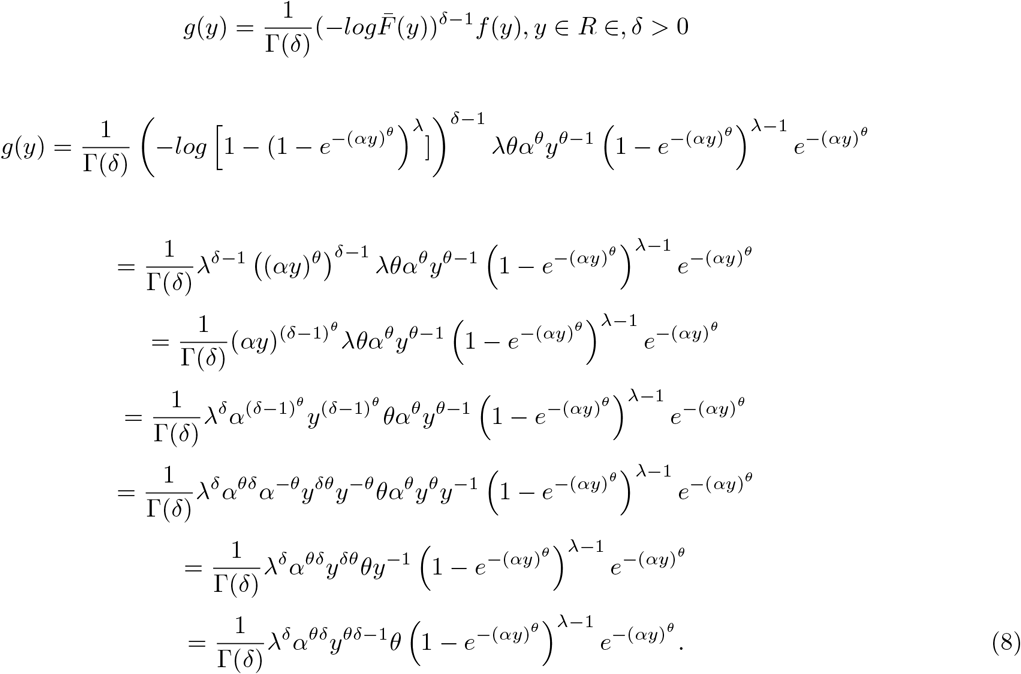

The Generalized Gamma Weibull (GGW) distribution stands out for its exceptional flexibility, as it encompasses several other sub-distributions as specific instances. This remarkable adaptability allows for an in-depth exploration of the diverse special cases inherent to the GGW distribution. Within the following subsections, we delve into some of the widely recognized special sub-models that fall under the Generalized Gamma Weibull distribution.

*Sub-model 1:*

If *λ* = 1 then the generalized gamma weibull distribution reduces to a generalized gamma distribution with parameter *δ, α, θ*, and its pdf is defined by [12]:

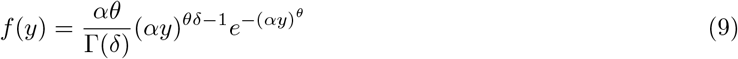

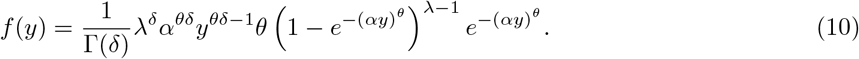

Substituting *λ* = 1 in equation 10 reduces to equation 9;

*Sub-model 2:*

If *λ* = 1 and *θ* = 1 then the generalized gamma weibull distribution reduces to gamma distribution and its pdf can be written as:

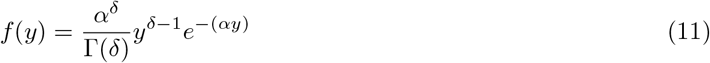

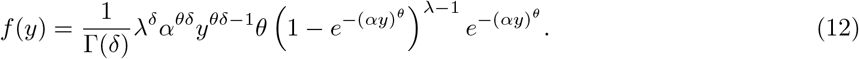

Replacing *λ* = *θ* = 1 in equation 12 reduces to equation 11.

*Sub-model 3:*

If *λ* = 1,*θ* = 1 and *δ* = 1, then the generalized gamma weibull distribution reduces to exponential distribution and its pdf can be written as:

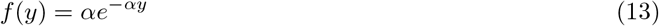

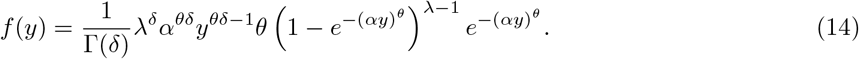

Replacing *λ* = *θ* = *δ* = 1 in equation 14 reduces to 13.

*Sub-model 4:*

If *δ* = 1 and *θ* = 1, then the generalized gamma weibull distribution reduces to Exponentiated Exponential distribution which is presented by [13] as follows:

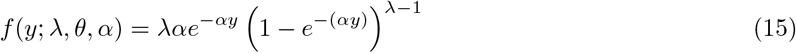

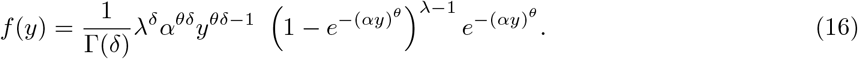

Substituting *δ* = 1 and *θ* = 1 in equation 16 reduces to equation 15.

The figures presented in Figure 1d display the (a) probability density function (pdf), (b) cumulative distribution function (Cdf), (c) hazard function, and (d) survival function of the new distribution. These plots exhibit varying shapes as the parameter values (*δ, θ, λ, α*) change. This variability is expected, as it aligns with the behavior of certain real-life datasets. Consequently, the modified distribution (GGW) demonstrates its capability to accommodate a wide range of shapes observed in real-world data.

**Figure 1:**
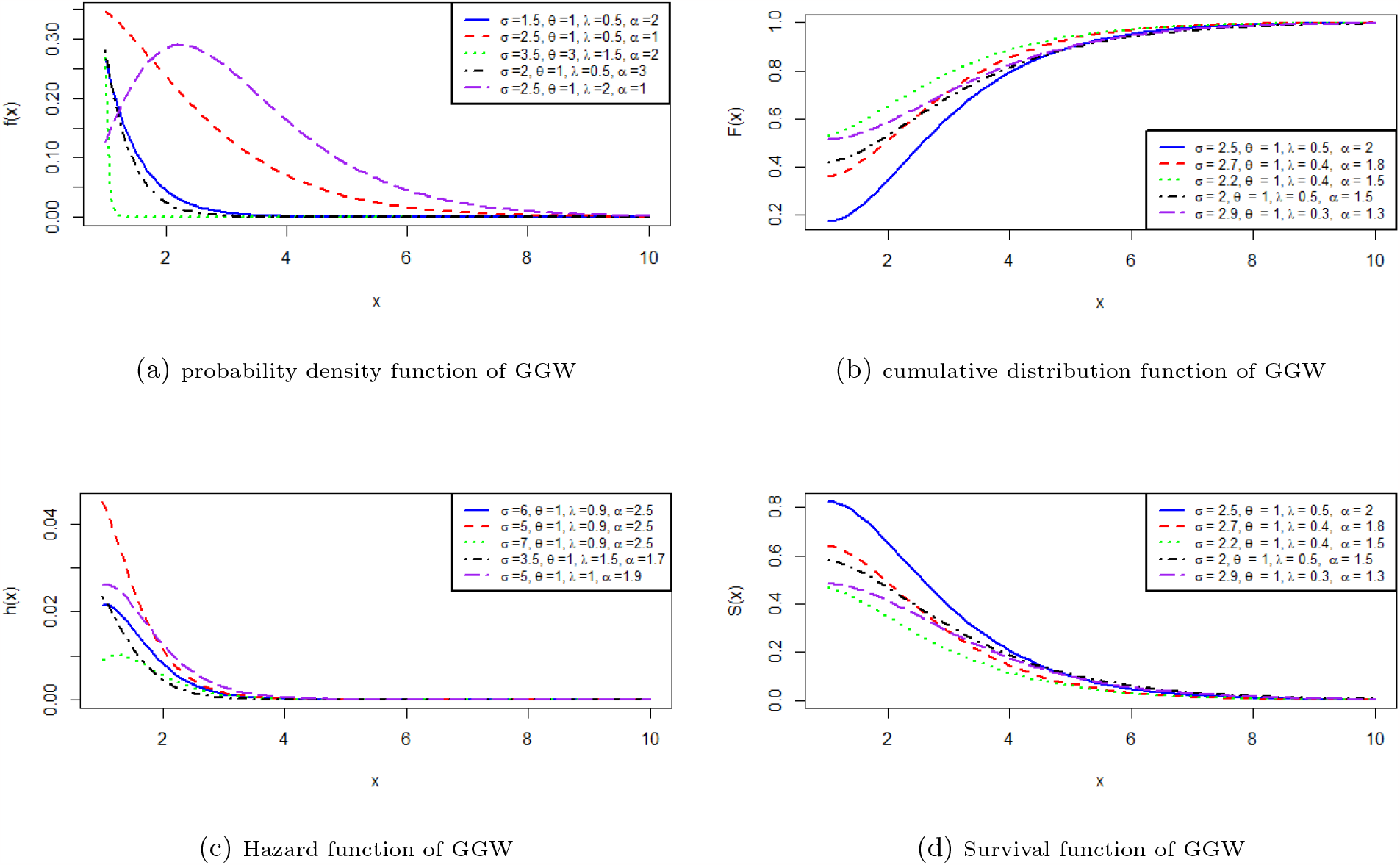
Plots of the four different functions of GGW

### 2.1 Hazard Rate Function (HRF)

The concepts of the Hazard Rate Function (HRF), sometimes referred to as the hazard function or instantaneous failure rate, are frequently employed in the disciplines of reliability engineering, survival analysis, and statistics[14].

Thus, the hazard rate function (HRF) for the Generalized Gamma Weibull distribution, as provided in equation 17, has been meticulously developed by precisely adhering to the quantity outlined in equation 3 and incorporating all the previously mentioned functions.

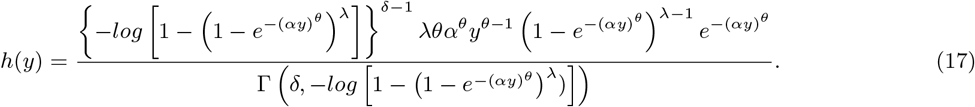

### 2.2 Expansion of the GGW Density Function

For any real parameter *δ >* 0,the following formula holds,

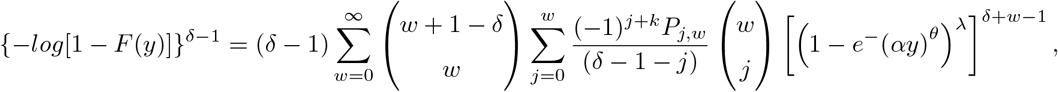

where *δ* is a real parameter and the constants *P*_*j,w*_ can be determine recursively by: 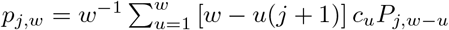 for *w* = 1, 2, … and *p*_*j*,0_ = 1. Furthermore, for any real parameter, we define:

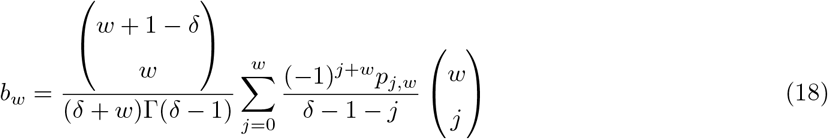

The corresponding equation 1 can be expressed as;

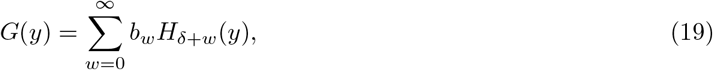

where *H*_*δ*+*w*_(*y*) denote the cdf of the exp(G)(*δ* + *w*) distribution. So, several properties of the gamma (G) distribution can be obtained by knowing those of the exp (G) distribution, see, for example, [15], [16] and [17], among others.

Then equation 5 can be expressed as equation 20 in other to find the linear combination of GGW distribution:

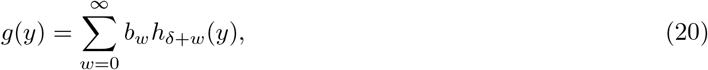

where *h*_*δ*+*w*_(*y*) denotes the pdf of the exp-G(*δ* + *w*) distribution. Then

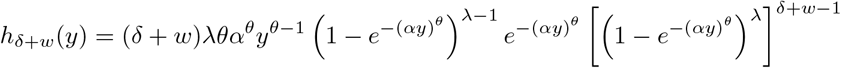

denotes the exponentiated generalized weibull (EGW) (*τ, δ* + *w*) density function. So, the GGW density function is a linear mixture of EGW density

### 2.3 Quantile Function of Generalized Gamma Weibull Distribution

In other to obtain the quantile function of the GGW, consider the use of the equation developed by gradshteyn[18].

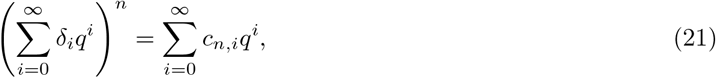

where the coefficients *c*_*n,i*_(*for*i=1,2…) are easily determined from the recurrence equation.

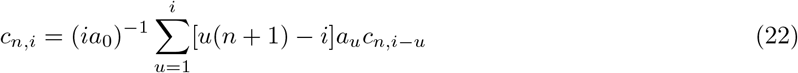

where 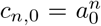. The coefficient *c*_*n,i*_ can be recursively calculated base on *c*_*n*,0_, …, *c*_*n,i*−1_. In mathematical terms, *c*_*n,i*_ can be expressed explicitly as a function of *a*_0_, …, *a*_*i*_, through this explicit expression might not be necessary for numerical programming using algebraic or numerical software.

Moreover, when considering a random variable y with a cumulative distribution function (CDF) denoted as F(y), we define the quantile of Y, denoted as (*y*_*q*_), as the inverse of the CDF, *F* ^−1^(*q*), where 0 *≤ q ≤* 1. This definition holds when substituting F(y) with q in equation 4. Consequently, the quantile function of the GGW distribution can be represented as follows:

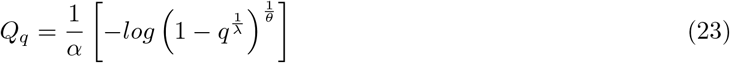

Let Y be a random variable following the GGW distribution. To generate random numbers from this distribution, we can utilize the expression in equation 23. Furthermore, the median of the GGW distribution, denoted as Med(Y), can be calculated by substituting q = 0.5 in equation 24 as follows.

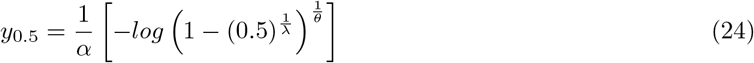

### 2.4 Moments of the GGW Distribution

Choudhury [19] developed a formula for calculating the jth raw moment of the EW distribution without imposing any constraints. This same approach was used to compute the expected value *E*(*Y*_*j*_). Additionally, the result *d*^*r*^(*e*^*ay*^)*/dy*^*r*^ = *a*^*r*^*e*^*ay*^ was implicitly utilized for 0 *≤ r ≤* 1, as demonstrated in prior research (example., [20])).

Consider transforming the generator in equation 2 using Laplace transform (LT) presented in equation 25 by letting x=F(z). Hence, the LT is then,

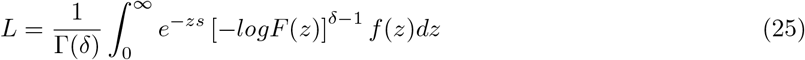

By defining x as a function of z, we get x = F(z).

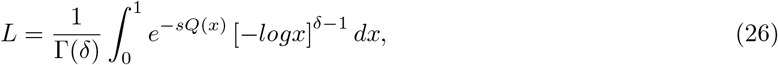

where *Q*(.) = *Q*_*GW*_ (.) indicates the quantile function of GW distribution. By expanding *e*^−*sQ*(*m*)^,

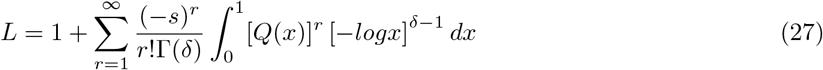

From F(y)=[1 *− e*^−(*αy*)^]^*λ*^, this integral reduces to

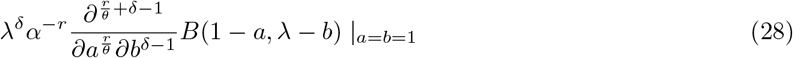

where 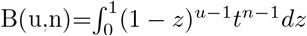 is the theta function. Hence,

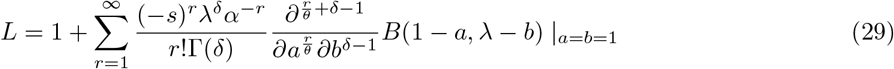

the *j*^*th*^ ordinary moment *μ*^′^_*j*_ follows as

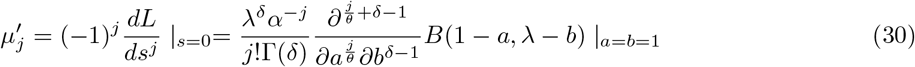

### 2.5 Order Statistics for the GGW Distribution

Order statistics play a vital role in various aspects of statistical theory and application. Consider a random sample *Y*_1_, *Y*_2_, …, *Y*_*n*_ drawn from a gamma distribution. Denote the *i*^*th*^ order statistic as *Y*_*i*:*n*_. These relationships can be further explored using Equation 19 and Equation 20.

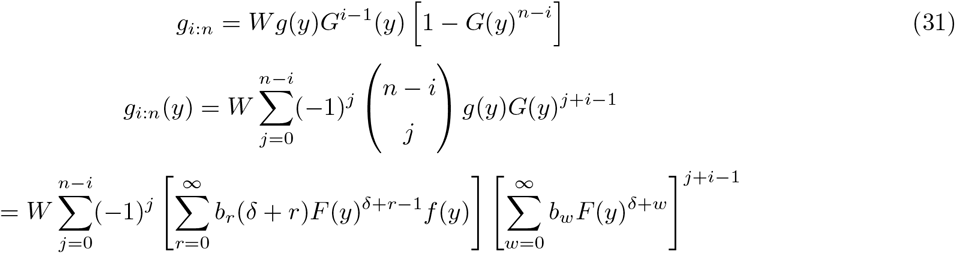

Where 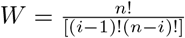 Using equation 21 and equation 22, we can write

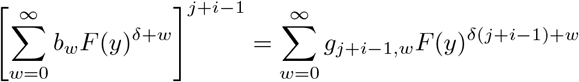

where 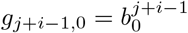 and,

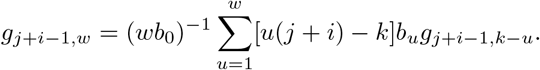

Hence,

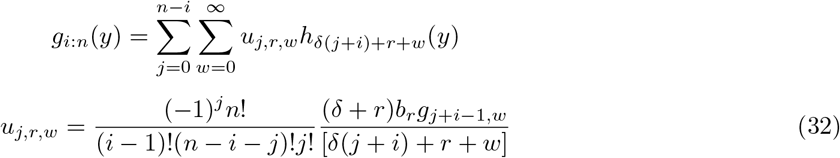

### 2.6 Maximum Likelihood Estimation for the GGW Distribution

Given a sample of size n, *y*_1_, …, *y*_*n*_, drawn from the GGW (*λ, θ, α, δ*) distribution, the log-likelihood function for the parameter vector Φ = (*λ, θ, α, δ*)^*T*^ is represented as follows:

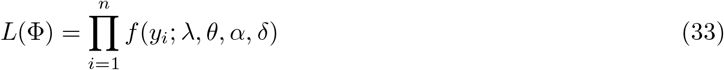

#### 2.6.1 Log-likelihood function for the GGW Distribution

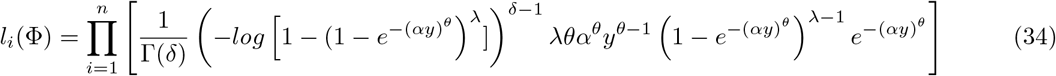

The log-likelihood function for Φ is

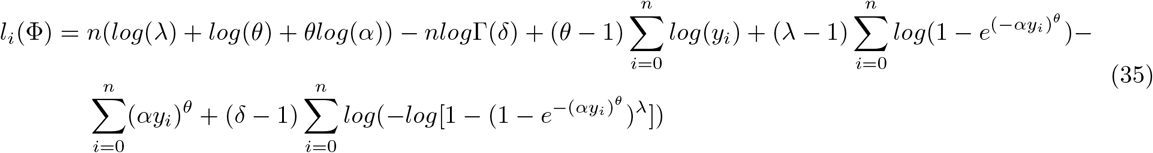

The partial derivatives of *l*(Φ) with respect to the parameters are

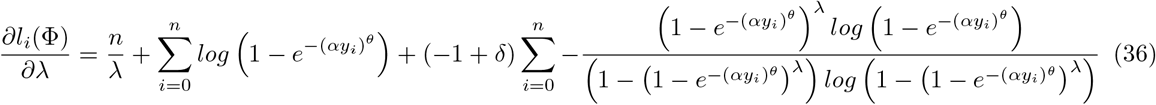

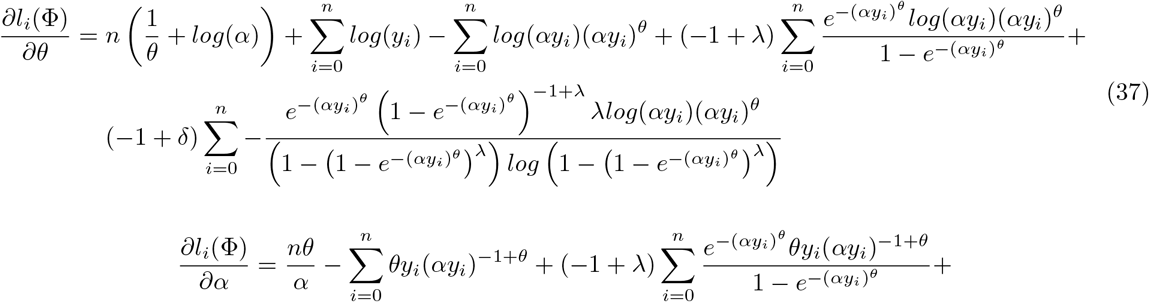

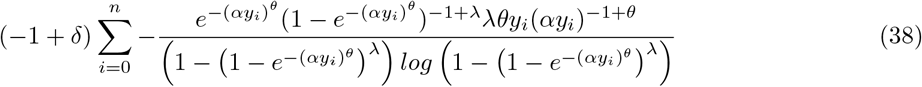

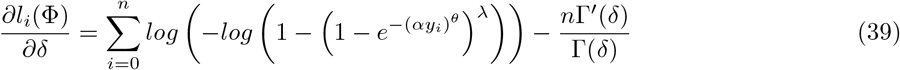

The maximum likelihood estimates (MLEs) for the parameters *λ, θ, α* and *δ*, denoted as 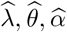 and 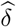 *δ* respectively, are determined by solving the following equations 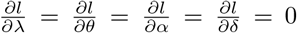 Since there is no closed-form solution for these equations, an optimization technique must be applied to find the optimal parameter values.

## 3. Asymptotic Confidence Intervals for the GGW Distribution

Numerical computation facilitates the derivation of expectations within the Fisher Information Matrix (FIM). Consider 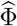, denoted as 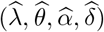 representing the maximum likelihood estimate of the parameter set Φ, which consists of (*λ, θ, α, δ*). Provided the standard regularity conditions hold and the parameters reside within the interior of the parameter space (without touching its boundary), we can establish that 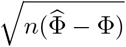 converges asymptotically to a normal distribution *N*_4_(0, *I*(Φ)^−1^), where *I*Φ represents the expected information matrix. To enhance clarity, we’ll replace *I*Φ with, 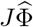 evaluated at 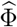, thus yielding a multivariate normal distribution *N*_4_(0, *I*(Φ)^−1^). In this distribution, the mean vector is 0 = (0, 0, 0, 0, )^*T*^ . This statistical framework enables the construction of confidence intervals and regions for various model parameters, including *λ, θ, α* and *δ*. These intervals are expressed as 100 (1 *− γ*) % confidence intervals. Furthermore, the subsequent section formulates and presents the maximum likelihood estimation with censored samples.

## 4. Maximum Likelihood Estimation in the Generalized Gamma Weibull Distribution with Censored Samples

Suppose a dataset with sample size ‘n’ consisting of independent positive random variables, denoted as *T*_1_, …, *T*_*n*_. Each *T*_*i*_ is associated with an indicator variable △_*i*_, which equals 0 if *T*_*i*_ represents a censoring time. Let’s define Φ as a parameter vector, Φ = (*λ, θ, α, δ*)^*T*^. In this context, the likelihood function is denoted as *L*(Φ), for a dataset that includes right-censored observations, represented as (*t*_1_, △_1_), …, (*t*_*n*_, △_*n*_), originating from a Generalized Gamma Weibull (GGW) distribution. The GGW distribution is characterized by its probability density function (pdf) and survival function, which would be employed to formulate the likelihood function.

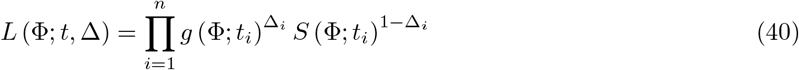

where *S*(*t*_*i*_) = 1 *− G*(*t*_*i*_).

The log-likelihood function, *l*(Φ), based on data, from equation 7 and equation 8 is

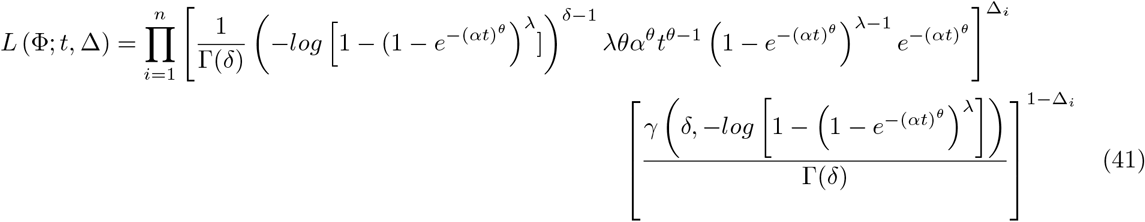

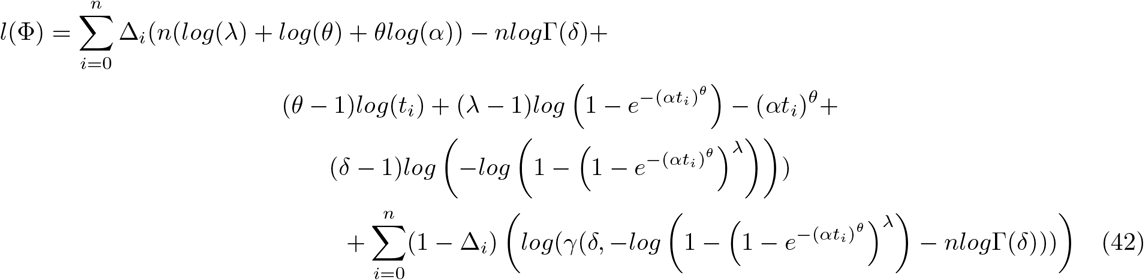

## 5. Methods of Evaluation

The aim of the real-life application is to select the best distribution when comparisons are made with respect to related existing distributions. To achieve this, the following criteria were used; Log-likelihood, Akaike Information Criteria (AIC), Bayes Information Criteria (BIC) and Corrected Akaika Information Criterion (CAIC).

AIC is defined as

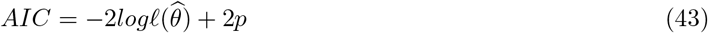

While BIC is defined as

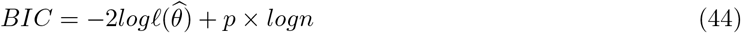

In the context of statistical modeling, 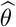 symbolizes the optimal parameter estimates derived from maximum likelihood estimation. 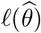 corresponds to the log-likelihood associated with these estimates. In both equation 43 and equation 44, the final components introduce a regularization term into the log-likelihood, which is contingent on the number of parameters involved. In linear models, these parameters are typically associated with regression coefficients. These regularization terms are employed to mitigate the risk of overfitting, and it’s worth noting that, for practical sample sizes, the regularization effect is more pronounced in the Bayesian Information Criterion (BIC) compared to the Akaike Information Criterion (AIC). Furthermore, for sufficiently large datasets, there exists the Corrected Akaika Information Criterion (CAIC).

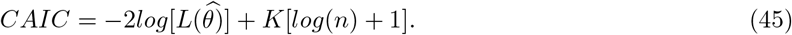

The theoretical framework of the Akaike Information Criterion (AIC) falls short when it comes to addressing situations in which the genuine model exhibits both significant effects and more gradual, diminishing influences (referenced as [21]). These particular aspects do not exhibit a direct correlation with the Kullback-Leibler discrepancy.

## 6. Application

The study delved into the application of the GGW distribution in the analysis of both censored and uncensored datasets. This exploration encompassed the examination and discussion of two real-life survival datasets and two real-life time datasets, constituting a total of four distinct datasets. The intent was to demonstrate that the GGW distribution holds promise as a robust lifetime model, particularly when juxtaposed with several well-known distributions, including but not limited to the Generalized Weibull, Generalized gamma, Gamma Weibull, Exponentiated Weibull, and Gamma distributions (referred to as G. Weibull, G. Gamma, Gamma W., E. Weibull, and Gamma, respectively). Furthermore, to gauge the performance of the GGW distribution, various statistical measures were employed, including the calculation of log-likelihood (L), Akaike information criterion (AIC), Corrected Akaike Information Criterion (CAIC), and Bayesian information criterion (BIC) values.

The statistical values under consideration encompass the Akaike Information Criterion (AIC), Corrected Akaike Information Criterion (CAIC), and Bayesian Information Criterion (BIC). Upon analyzing these values, it becomes evident that the GGW model demonstrates a superior fit to the provided data.

Table 1 provides estimates of the GGW distribution parameters and its related sub-models, accompanied by standard errors in parentheses. Additionally, it provides the AIC, CAIC, and BIC values for the Aarset dataset. Furthermore, the estimated covariance matrix for the GGW model can be shown below:

**Table 1:** GGW Estimation for Life time of 50 devices ([22]).

| Distribution | Estimate |  |  |  | Statistic |  |  |  |
| --- | --- | --- | --- | --- | --- | --- | --- | --- |
| | $\lambda$ | $\theta$ | $\alpha$ | $\delta$ | -2 log Likelihood | AIC | BIC | CAIC |
| **GGW** | 0.6316 | 0.0777 | 0.0037 | 0.3286 | 330.6313 | **338.6313** | **346.2794** | **339.5202** |
| **G Weibull** | 1.5438 | 0.7646 | 0.0090 |  | 370.1946 | 376.1946 | 381.9307 | 376.5424 |
| **Gamma Weibull** |  | 2.435 | 0.650 | 0.843 | 387.2605 | 393.2605 | 398.9965 | 393.7822 |
| **E W** | 6.360 | 0.690 | 3.205 |  | 411.0439 | 417.0439 | 422.7800 | 417.5657 |
| **Gamma** |  |  | 0.0175 | 0.8002 | 480.3805 | 484.3805 | 488.2045 | 484.6358 |
| **G Gamma** |  | 0.500 | 1.1466 | 1.462 | 1040.847 | 1046.847 | 1052.583 | 1047.369 |

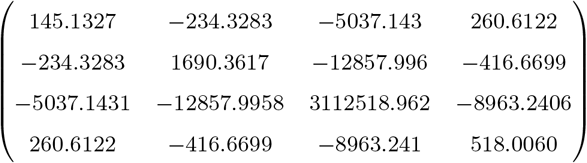

The 95% asymptotic confidence intervals are: *λ ϵ* 0.6316 *±* 1.96(12.0471), *θ ϵ* 0.0777 *±* 1.96(41.1140), *α ϵ* 0.0037 *±* 1.96(1764.2333) and *δ ϵ* 0.3286 *±* 1.96(22.7598). The second example consists of lifetime data relating to relief times (in minutes) of patients receiving an analgesic.

Table 2 presents parameter estimates for the GGW distribution and its associated sub-models, as well as the AIC, CAIC, and BIC values for lifetime data pertaining to the relief times (measured in minutes) of patients administered an analgesic [23]. Additionally, the estimated covariance matrix for the GGW model is given by:

**Table 2:** GGW Estimation for lifetime data relating to relief times (in minutes) of patients receiving an analgesic.

| Distribution | Estimate |  |  |  | Statistic |  |  |  |
| --- | --- | --- | --- | --- | --- | --- | --- | --- |
| | $\lambda$ | $\theta$ | $\alpha$ | $\delta$ | -2 log Likelihood | AIC | BIC | CAIC |
| **GGW** | 0.6027 | 0.0789 | 0.0450 | 0.3446 | 33.9009 | **41.9009** | **45.8839** | **44.5676** |
| **Gamma** |  |  | 2.1589 | 4.1020 | 41.5398 | 45.5398 | 47.5312 | 46.2457 |
| **G Weibull** | 2.3000 | 2.4200 | 0.2967 |  | 66.2335 | 72.2335 | 75.2207 | 73.7335 |
| **Gamma W** |  | 2.4350 | 0.6500 | 1.8209 | 83.0377 | 89.0377 | 92.0249 | 90.5377 |
| **E Weibull** | 6.3600 | 2.7612 | 12.3400 |  | 282.8704 | 288.8704 | 291.8576 | 290.3704 |
| **G Gamma** |  | 2.2500 | 0.3461 | 1.4616 | 447.8106 | 453.8106 | 456.7978 | 455.3106 |

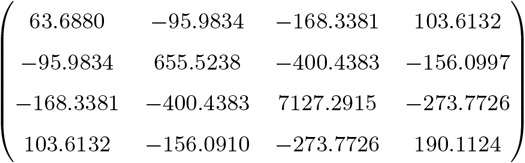

The 95% asymptotic confidence intervals are: *λ ϵ* 0.6027 *±* 1.96(7.9805), *θ ϵ* 0.0789 *±* 1.96(25.6032), *α ϵ* 0.0450 *±* 1.96(84.4233) and *δ ϵ* 0.3446 *±* 1.96(13.7881)

### 6.1 Long-term survival with random censoring

The incorporation of long-term survival considerations necessitates adjustments to both the probability density function (PDF) and the survival function. To address this, two distinct datasets were analyzed: one from the Veterans’ Administration Lung Cancer study [24] and the other from the Statistical Analysis of Failure Time Data and North Central Cancer Treatment Group for Lung Cancer [25]. Table 3 and Table 4 present estimates of various parameters for right-censored models, alongside metrics such as AIC (Akaike Information Criterion), CAIC (Consistent Akaike Information Criterion), and BIC (Bayesian Information Criterion).

**Table 3:**
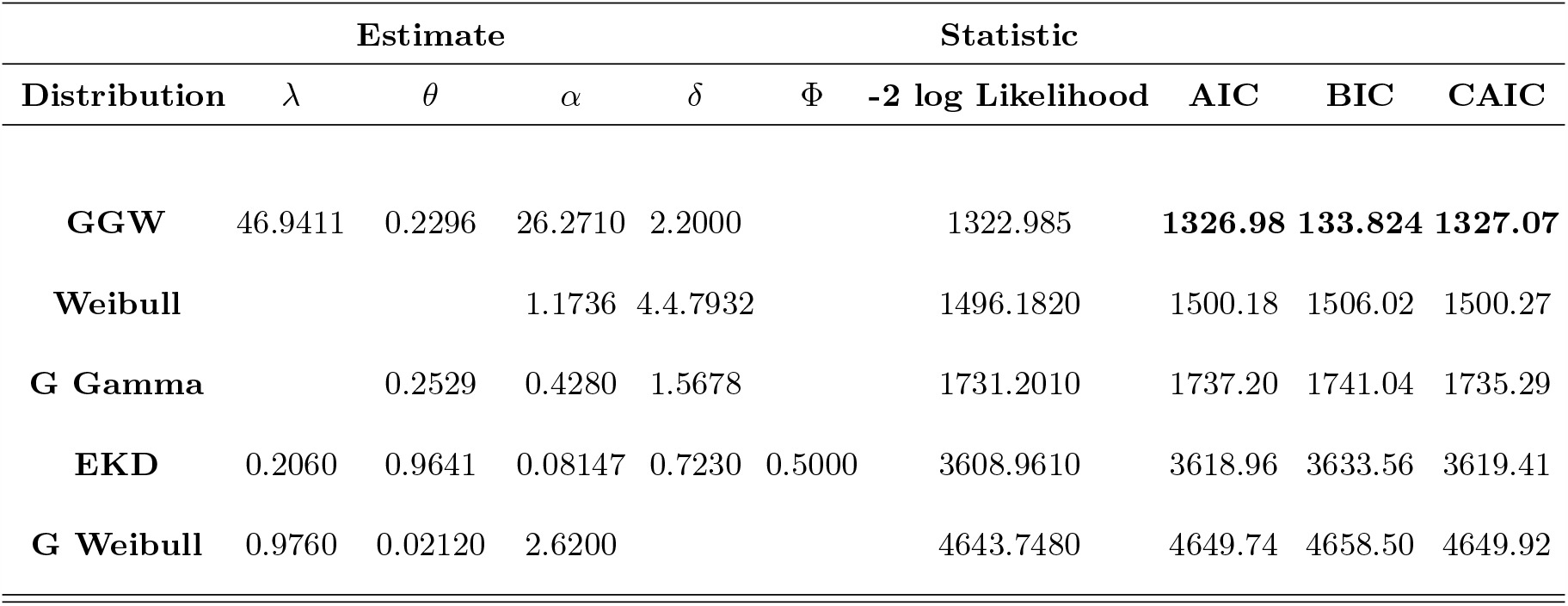
GGW Estimation for Veterans’ Administration Lung Cancer study.

**Table 4:** GGW Estimation for North Central Cancer Treatment Group for Lung Cancer.

| Distribution | Estimate |  |  |  | Statistic |  |  |  |  |
| --- | --- | --- | --- | --- | --- | --- | --- | --- | --- |
| | $\lambda$ | $\theta$ | $\alpha$ | $\delta$ | $\Phi$ | -2 log Likelihood | AIC | BIC | CAIC |
| **GGW** | 0.9285 | 0.2000 | 19.2000 | 5.2000 |  | 4824.7575 | **4832.757** | **4846.474** | **4832.936** |
| **Weibull** |  |  | 1.0900 | 5.1085 |  | 4885.9930 | 4891.993 | 4891.140 | 4896.263 |
| **G Gamma** |  | 0.1078 | 0.4280 | 1.3599 |  | 6845.0630 | 6851.063 | 6861.351 | 6851.170 |
| **EKD** | 0.0504 | 0.0018 | 0.0450 | 0.0049 | 0.0062 | 6985.0832 | 6995.084 | 7012.231 | 7000.135 |
| **G Weibull** | 4.8000 | 0.2500 | 5.7600 |  |  | 7841.1200 | 7841.1800 | 7841.2830 | 7841.1181 |

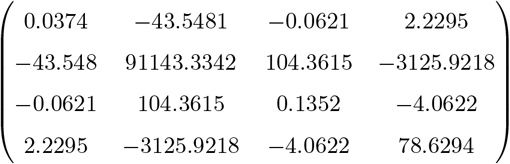

The 95% asymptotic confidence intervals are: *λ ϵ* 46.9411 *±* 1.96(0.1934), *θ ϵ* 0.2296 *±* 1.96(301.8910), *α ϵ* 26.2710 *±* 1.96(0.3677) and *δ ϵ* 2.2000 *±* 1.96(8.8673)

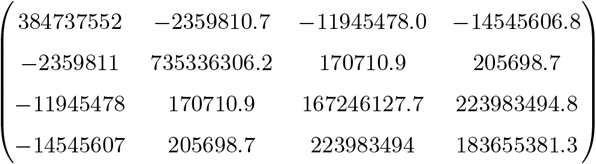

The 95% asymptotic confidence intervals are: *λ ϵ* 0.9285 *±* 1.96(19614.728), *θ ϵ* 0.2000 *±* 1.96(27117.085), *α ϵ* 19.2000 *±* 1.96(12932.367) and *δ ϵ* 5.2000 *±* 1.96(13551.951)

The GGW distribution emerges as a formidable contender when juxtaposed with its counterparts featuring three and four parameters distributions, as indicated in Tables 1-4. In this evaluation, we rely on key values such as the AIC, BIC, and CAIC. Remarkably, the GGW distribution consistently yields the most favorable results, exhibiting the lowest values for AIC, BIC, and CAIC across both censored and uncensored datasets. When assessing model performance based on these information criteria, the clear preference leans towards selecting the model with the lowest information criteria value.

## 7. Conclusions

This study introduces a new modified distribution known as GGW, which represents a significant advancement in statistical modeling. Extensive efforts have been dedicated to exploring and substantiating the estimations and properties associated with this new modified distribution. Additionally, the practical applicability of GGW has been demonstrated using real-life data in the R programming environment. Notably, the outcomes of this investigation showcase the superior performance of GGW in comparison to existing distributions, as evidenced by the AIC, BIC, and CAIC criteria.

Furthermore, the Generalized Gamma Weibull (GGW) distribution, an encompassing framework that subsumes the generalized gamma Weibull, gamma, exponential, and exponentiated exponential distributions as special cases, is proposed. This comprehensive approach delves into diverse aspects of GGW, including moments, the quantile function, median, and order statistics, with a focus on deriving these properties. The parameter estimation for GGW is accomplished through the maximum likelihood method, and its efficacy is scrutinized across varying sample sizes and parameter combinations. The overarching characteristics of the new modified distribution can be revealed by representing its density function as a linear combination of gamma exponentiated Weibull density functions. Additionally, alternative expressions for moments have been derived using fractional derivatives, an aspect that often goes overlooked in statistical studies, despite its widespread utility in Engineering and Physics. As an illustrative example, we refer to Lemma 5 (b) from Zografos and Balakrishnan [1], which pertains to gamma-F distribution and emphasizes the relevance of these derivatives in statistical analysis. It states that

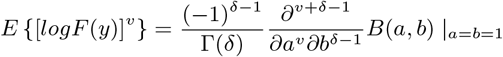

The condition of natural *δ >* 1 and the presence of variable v lead to a significant outcome. This result remains consistent for any positive real value of v, achieved through the utilization of fractional derivatives. It is highly probable that these derivatives can be applied to compute raw moments for various distributions belonging to the gamma (G) and gamma 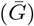 classes.

for natural *δ >* 1 and *v*. This result holds for any positive real *v* by using fractional derivatives. It is very likely that these derivatives can be used for many members of the gamma (G) and gamma 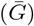 classes of distributions as a way of obtaining their raw moments.

